# Improving the Power of Structural Variation Detection by Augmenting the Reference

**DOI:** 10.1101/019109

**Authors:** Jan Schröder, Santhosh Girirajan, Anthony T. Papenfuss, Paul Medvedev

## Abstract

The uses of the Genome Reference Consortium’s human reference sequence can be roughly categorized into three related but distinct categories: as a representative species genome, as a coordinate system for identifying variants, and as an alignment reference for variation detection algorithms. However, the use of this reference sequence as simultaneously a representative species genome and as an alignment reference leads to unnecessary artifacts for structural variation detection algorithms and limits their accuracy. We show how decoupling these two references and developing a separate alignment reference can significantly improve the accuracy of structural variation detection, lead to improved genotyping of disease related genes, and decrease the cost of studying polymorphism in a population.

## 1 Introduction

The initial sequencing and assembly of a human reference genome allowed for the understanding of our genomic landscape in comparison to other species [1, 2]. It also facilitated our understanding of polymorphism within the human species by providing a high-resolution coordinate system onto which variants could be mapped [2]. As resequencing projects became wide-spread, the reference also began to play a central role as a tool for variant detection and discovery algorithms. By mapping the reads to the reference, one could identify both structural and single-nucleotide variants in the sequenced (donor) genome.

Thus, the uses of the human reference sequence can be roughly categorized into three related but distinct categories: as a representative species genome, as a coordinate system for identifying variants, and as an alignment reference for variation detection algorithms. The reference sequence used for all the above scenarios is maintained by the Genomic Reference Consortium (GRC). One notable exception is the idea of a human pan-genome, which has been introduced [3] to distinguish the representative species genome from the GRC reference.

The use of the GRC reference genome as an alignment reference has led to some artifacts in the structural variants we can detect. One striking example is that most structural variation (SV) detection methods have less power to detect long insertions than deletions, with respect to the GRC reference [4]. Identifying large insertions is notoriously difficult, since it requires careful de novo assembly procedures and the detection of two novel adjacencies [5, 6]. Deletions, on the other hand, are significantly easier, since only one new adjacency has to be detected and no novel sequence has to be considered [7]. However, whether an indel is a deletion or insertion depends on which allele sequence is represented in the GRC reference genome. Thus the power to detect a variant depends on the sequence content of the GRC reference. Such artifacts seem unnecessary and arbitrary and can pose challenges to downstream analyses, as large indel polymorphisms play a key role in the susceptibility to disease of individuals or entire populations.

We propose that the alignment reference should be decoupled from the traditional GRC reference. The alignment reference can be considered as simply a sequence of nucleotides that serve as an input to variant detection algorithms, as opposed to a representative genome or a coordinate system for mapping variants. This sequence does not need to represent a real or even mosaic genome. We can then pose the question: what sequence would maximize the power of SV detection algorithms?

In this paper, we demonstrate how a distinct alignment reference genome can be used to increase the power to detect insertions. First, we show how to construct an alignment reference by augmenting the GRC reference with known insertions. We use a set of insertions found in the HuRef genome [8], relative to the GRC reference. We then develop a pipeline that “wraps” around any existing SV calling pipeline to incorporate the augmented reference. Finally, we run this pipeline on low-coverage sequencing data from 16 individuals from the 1000 Genomes Project and show that the accuracy of detecting the insertions increases by 67%.

## 2 Results

We first identified 229 high confidence insertions in the HuRef genome (Supplementary Fig. 1), which is an alternative human whole genome assembly based on 454 sequencing data from J. Craig Venter [8]. These are insertions in HuRef relative to the GRC reference (hg18) that are at least 300nt in length and do not lie within 300nt of a repetitive region. We refer to those as Venter Novel Alleles (VNAs). We then created an augmented alignment reference, called ref+, by injecting the sequence of the VNAs into the appropriate locations of hg18. Ref+ contains 328kbp of new sequence, covering 48 genes. We note that none of the VNAs are present in the database of genomic structural variation (dbVar), except as entries from the HuRef study itself.

**Figure 1:**
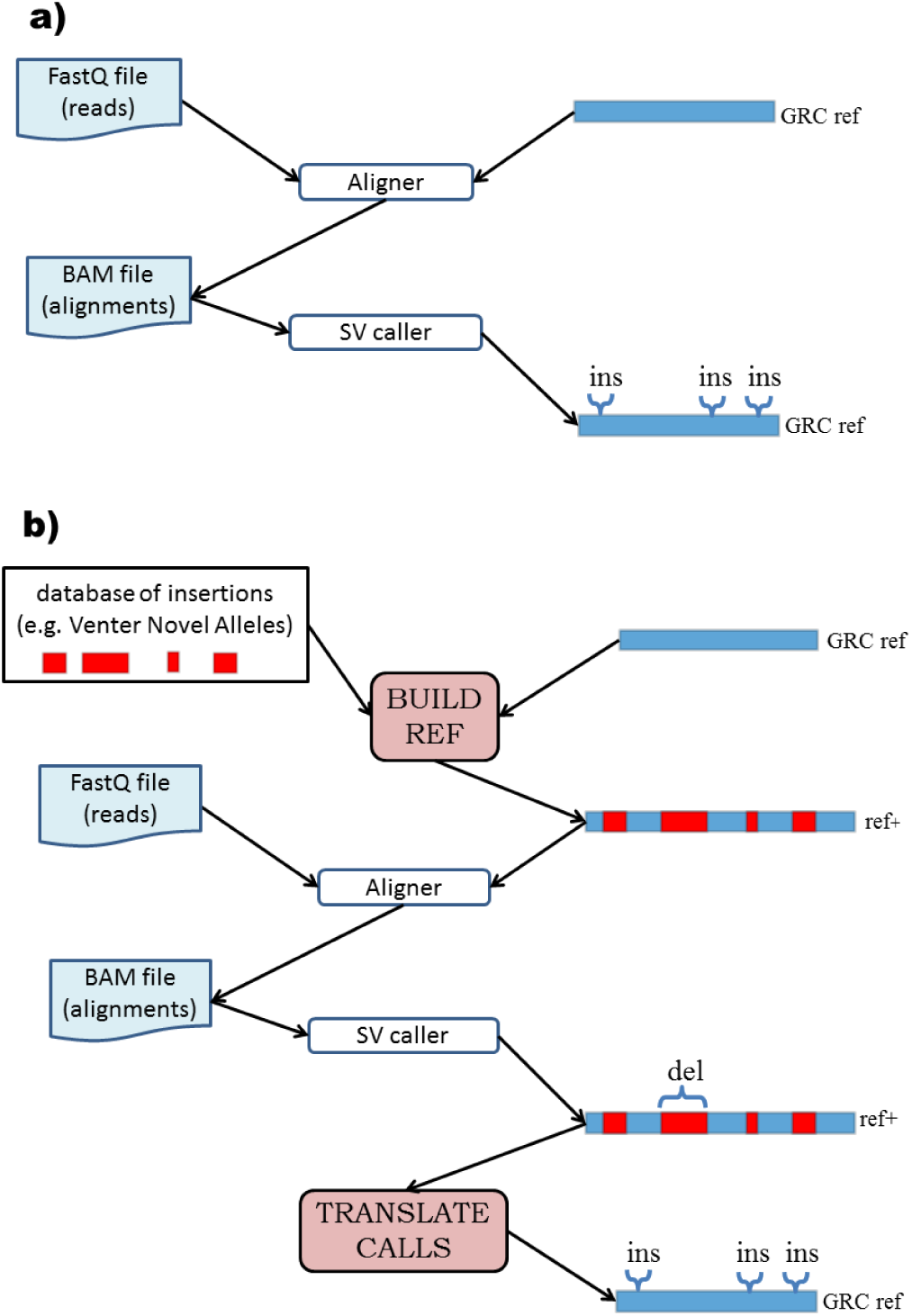
Method workflow. **a)** In a traditional SV calling pipeline the reads are first aligned against the GRC reference and the alignments are passed to an SV caller, which annotates regions of the GRC reference as being inserted/deleted. **b)** Our approach is composed of two additional components. BUILD_REF takes a set of sequences to be inserted and modifies the GRC reference genome (e.g. hg18) by inserting the sequences into their prescribed locations, obtaining a new genome (ref+). We next align the reads to ref+ and run a SV caller. The TRANSLATE_CALLS component then modifies the resulting calls so that they become calls relative to the GRC reference, not ref+.

A typical SV detection pipeline maps the reads to the GRC reference genome, runs an SV caller to analyze the resulting mappings for SV signatures, and then outputs a set of loci in the GRC reference that are the location of the called SVs (Fig. 1a). We demonstrate how to modify any such pipeline to use ref+ instead (Fig. 1b). After creating ref+, we align the reads to ref+ and run the SV caller. The SV caller now reports calls relative to ref+, so we convert these to be relative to the GRC reference: deletions in injected regions correspond to no variation relative to the GRC reference, while no-calls in injected regions correspond to insertions relative to the GRC reference (see Methods section for more details). The potential power of using ref+ instead of the GRC reference is illustrated in Figure 2.

**Figure 2.**
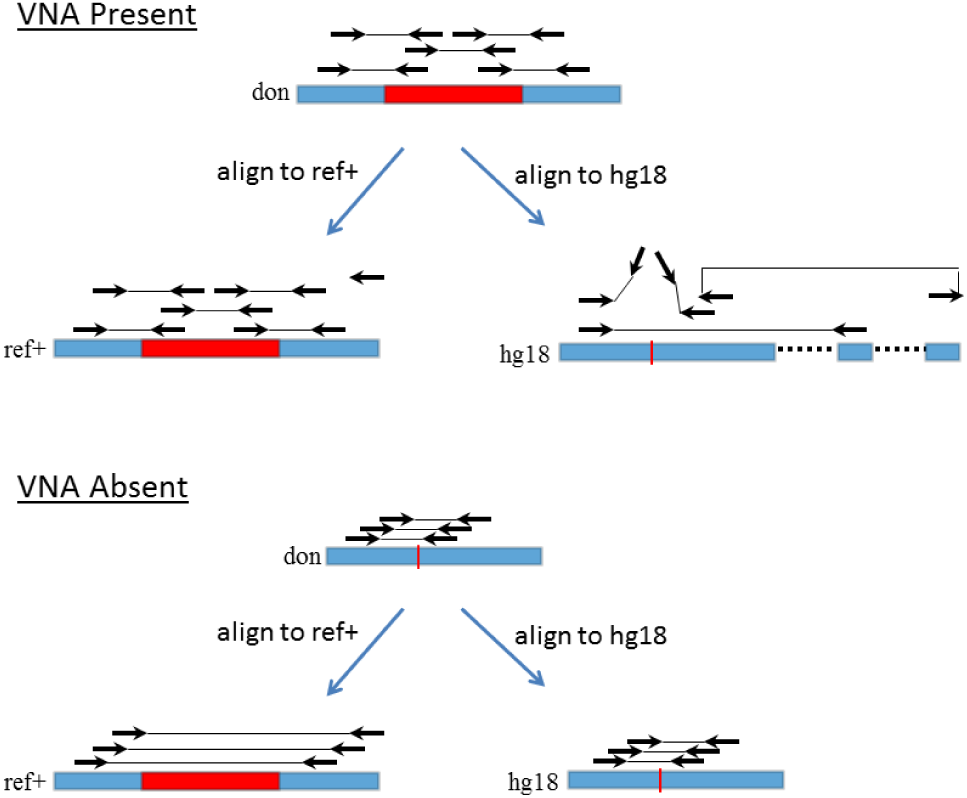
An illustrative example. In the top scenario, a VNA (shown in red) is present in the donor. In ref+, only concordant alignments (correct orientation and mapped distance) are present. As a result, the SV caller does not make a call in ref+, which is converted by TRANSLATE_CALLS to an insertion call in the GRC reference (hg18). In the GRC reference, however, the read pairs that originate from across the VNA junction map discordantly, with one read left unmapped or falsely mapping to a homologous region. These signals in the GRC reference are difficult to decipher for any SV algorithm. In the bottom scenario, where the VNA is absent in the donor, the pairs that span the VNA injection point in the donor align concordantly to the GRC reference. In ref+, they align discordantly with an enlarged mapped distance but bear the hallmark signature of a deletion. This is among the easiest signals that an SV caller can detect and most algorithms show good results with respect to this SV type

We wanted to demonstrate the power of using ref+ with existing pipelines to detect SVs in a population setting of multi sample, low coverage sequencing data. We used 1000 Genomes Project [9] data for 16 individuals, with five individuals each from the YRI and CHB populations and six individuals from the CEU population (Supplementary Table 1). We used bowtie2 [10] as the aligner and Delly [11] as the SV caller, which are common tools used for SV detection. We ran both the standard GRC pipeline and the ref+ pipeline (raw results in Supplementary Table 2), and measured the accuracy as the proportion of validated sites that were correct (see Methods section for validation details).

The average accuracy of the ref+ pipeline was 80% (σ=5%) while the accuracy of the GRC pipeline was 48% (σ=13%), an increase of 67% (Fig. 3, Supplementary Table 3). As expected, the GRC pipeline had low sensitivity (average of 8.3%) compared to the ref+ pipeline (77.7%). The false discovery rate (FDR) was higher with the ref+ pipeline (30.6% average) than with the GRC pipeline (16.0% average), since the GRC pipeline made much fewer calls (avg=12) than did the ref+ pipeline (avg=96). However, the increase in sensitivity outweighed the decrease in FDR, as the average increase in the accuracy per sample was 31 percentage points.

**Figure 3.**
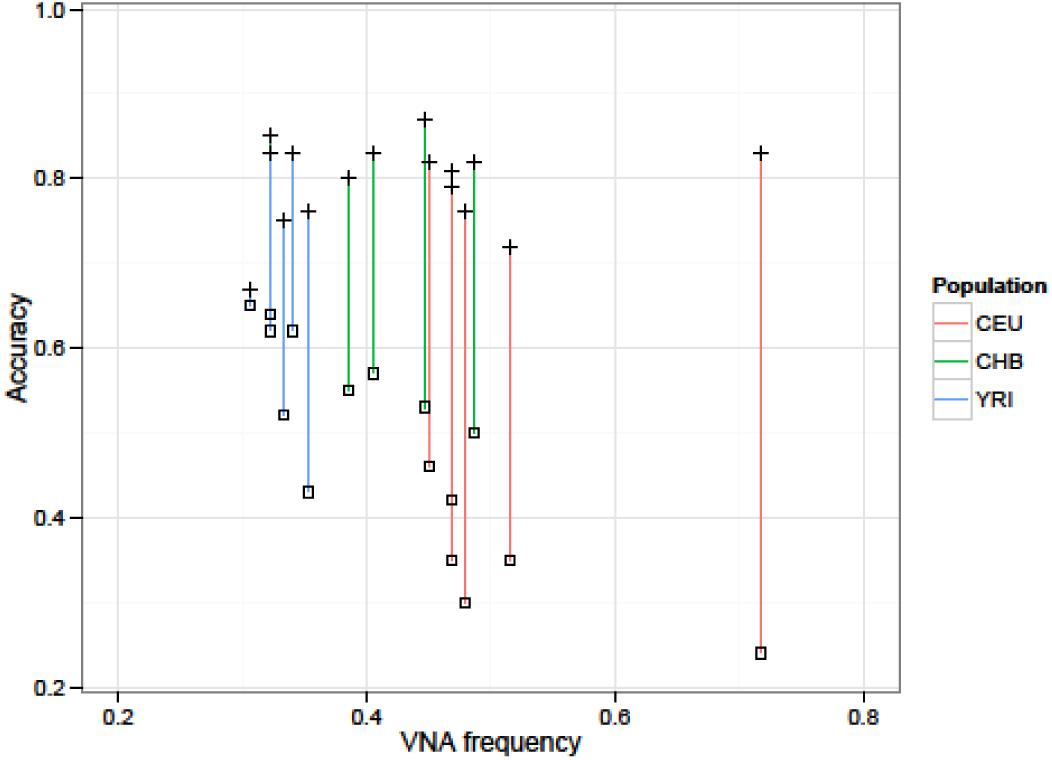
Analysis of ref+ pipeline accuracy. Each vertical line represents one individual, with the plus (+) point representing the ref+ pipeline and the square point representing the GRC pipeline.

We observe that the increase in accuracy depends on the population. In the CEU population, the mean increase was 44 percentage points, while in the CHB and YRI populations it was 28 and 20 points, respectively (Supplementary Table 3). This is expected, given that we found that VNAs are more frequent in CEU than in CHB, and more frequent in CHB than in YRI (Supplementary Fig. 2). These findings for VNAs are consistent with the known genetic heritage of the HuRef genome [8].

## 3 Discussion

The increased power offered by ref+ can help in genotyping variants of clinical importance, as some of the VNAs affect genes that play a role in disease. For example, our ref+ pipeline was able to detect an ALU insertion in the intronic region of CNTNAP2, a gene associated with autism and schizophrenia (the GRC pipeline did not detect this insertion). In general, an augmented reference can be used to target any known insertions of special interest. The approach here can be extended to include other novel sequences, such as the alternate haplotypes included with GRCh38.

Our results also suggest that an augmented reference can be used to decrease the costs of polymorphism discovery and detection in a population study. A single genome can be sequenced at high coverage to allow to de novo assemble novel insertions using methods such as [12, 5] and include them in an augmented reference. Other individuals in the population can then be sequenced at low coverage while allowing the detection of the novel insertions. A similar approach can be applied in a family setting, by sequencing the parents at high and the children at low depth.

The use of ref+ is not always recommended. For instance, if the goal is to detect SNPs, then the presence of repetitive VNAs in the alignment reference may create false mappings, thus decreasing SNP detection accuracy. Or, for detecting SVs in populations that are expected to have a low frequency of VNAs, the higher false discovery rate may outweigh the benefits of better sensitivity.

The availability of longer reads (e.g. PacBio) can simplify the task of detecting variants longer than the Illumina read lengths. However, many insertions will remain too large to be captured even by long reads. Moreover, the use of long read technologies is still limited, the impact of their different error properties is yet to be fully assessed, and it is not clear if their use will become ubiquitous or limited to certain applications.

We have argued for the need to decouple the reference used for alignment from the reference used as a representative species genome. Our results indicate just one possible way that an alignment reference can be constructed to improve SV detection. Undoubtedly, the development of new ideas will lead to approaches that improve accuracy even further. Ideally an alignment reference would capture all the possible alleles by using a graph, but such an approach would require more sophisticated alignment algorithms. In fact, two recent papers have shown how reads can be efficiently and accurately aligned to a reference graph that contains multiple genomes from a population [13, 14]. The further development of such graph alignment algorithms will enable more sophisticated approaches to building the best alignment reference. However, the trade-offs involved between representing a more complete set of alleles (e.g. graph based approach) and allowing the use of existing alignment methods (e.g. linear based approach such as ref+) are not yet clear.

## 4 Methods

### 4.1 Data

We identified 229 Venter Novel Alleles (VNAs) as the insertions in HuRef [8] that meet the following criteria: the insertion locus (the locus in between two nucleotides in the GRC reference) does not fall into a repeat (according to the RepeatMasker track from the UCSC genome browser), is not within 300nt of a tandem repeat (simpleRepeat track on the UCSC genome browser), has unique mappability (100% according to the wgEncodeCrgMapabilityAlign100mer track on the UCSC genome browser), and the inserted sequence has a length greater than 300nt. Supplementary Figure 1 visualizes these using the PhenoGram software [15]. These filters are intended to select a set of alleles which have the best potential to be detected with Illumina sequencing. Alleles that are embedded in repeats would be difficult to detect for both the GRC and ref+ pipelines, interfering with the interpretation of our results. Alleles shorter than 300nt are below Delly’s detection threshold on the available data.

We selected 16 individuals from the 1000 Genomes Project as testing data (six with European background (CEU), five Chinese (CHB), and five African (YRI)) (Supplementary Table 1). We chose individuals to achieve a balance of background and to avoid related individuals (i.e. trios). We also chose the individuals so that we had a high coverage of libraries with at least 100nt reads and consistent insert sizes (around 350-450nt). We used only such runs since Delly utilizes a combination of split-read and paired-end information in the data to generate its output, and is therefore dependent on long reads as well as consistent insert sizes.

### 4.2 Ref+ construction

The augmented reference ref+ is constructed by creating new chromosomes that inject VNAs into the specified coordinates of hg18 (the build_ref script, Fig. 1b). This increases the chromosome sizes and coordinates shift towards higher positions. We therefore generate a set of offsets that allows coordinate transfer between ref+ and hg18. The translate_calls script uses these offsets to translate calls relative to ref+ into calls relative to hg18. Calls in non-injected regions are simply converted onto the corresponding co-ordinates of the hg18 reference. Deletion calls in injected regions correspond to no variation relative to the GRC reference, while no calls in injected regions correspond to insertions relative to the GRC reference.

### 4.3 SV calling pipeline

To analyse the impact of different reference genomes, we create a standard bioinformatics pipeline that can be used in any project that analyzes SVs in NGS data. We chose a single algorithm to perform the task of variant detection: Delly [11]. We chose Delly because it offers dedicated modules for deletion and duplication detection, and has been used in large-scale SV analyses [16]. However, any SV detection tool could be used. Reads are mapped to the reference genome (ref+ in the ref+ pipeline and hg18 in the GRC pipeline) with bowtie2 (2.0.0 beta 7) in local mode. Then, Delly sub-modules are executed on aligned reads (delly for ref+, duppy for hg18). Delly version 0.0.9 is used. Next, the set of SV calls from Delly are analysed with respect to the VNA sites. In hg18, duplications called within 500nt of a VNA insertion sites are regarded as predictions of VNA insertion. In ref+, we compare the deletion calls to the intervals corresponding to the VNA sites, and establish a Delly deletion of the VNA if the intervals overlap with an F-score higher than 0.1. The F-score is defined as *2PR/(P+R)*, where R is the proportion of the VNA covered by a Delly call (recall) and P is the proportion of the respective Delly call inside the VNA (precision). Finally, ref+ calls are translated into hg18 calls using the translate_calls script described above.

### 4.4 Accuracy calculation

We establish the accuracy of the ref+ and hg18 pipelines on account of how well they agree with the validation classifier. The validation classifier is our independent method to establish the allele status at a particular site and is described in the next section (Sec. 4.5). For each VNA site where the validation classifier is able to establish the status of the donor allele, we categorize it as a true positive (TP) or negative (TN) if our pipeline call agrees with the validation, or as a false positive (FP) or negative (FN) otherwise. More specifically, a TN is accounted for in hg18 if Delly does not call the site and the classifier evaluated the reference allele to be present homozygously; a homozygous reference allele paired with an insertion call by Delly is considered a FP; if both the alleles are present (heterozygous state) or the VNA is present homozygously, but Delly does not make call, it is a FN, otherwise a TP. Analogously, Delly deletion calls in ref+ are evaluated as TP for homozygous and heterozygous reference alleles, absent calls as FN; for homozygous VNAs a Delly call means a FP, and a TN upon absence of a call. The contingency tables for each of the samples for hg18 and ref+ are shown in Supplementary Table 3. We use the standard formulas to calculate the accuracy as *(TP+TN)/(TP+TN+FP+FN)*, the sensitivity as *TP/(TP+FN)*, and the false discovery rate as *FP/(TP+FP)*.

### 4.5 Validation classifier details

We designed our own classifier to assess VNAs upon their presence or absence in the samples, independently from the SV calling pipeline. The purpose of this classifier is to establish the true status of each VNA in a sample, so that we can evaluate the performance of the SV pipeline. The classifier operates with the knowledge of the VNA’s loci, and joins the signal from reads mapped to hg18 as well as ref+. Additionally, the sequencing data used by the classifier is a superset of that available to Delly: some, but not all, of the samples have runs with different library preparation available to them. Delly needs a homogeneous distribution of fragment lengths, but our classifier makes use of all the runs available. The read coverage utilised to classify alleles in each individual as well as the run accession numbers is shown in Supplementary Table 4.

The classifier establishes evidence for the reference allele if there are at least three reads spanning the VNA insertion site in hg18. We define a read as spanning if it overlaps the locus by at least 10nt on either side (this requirement is designed to exclude mis-mapped and soft-clipped reads from the classification). The classifier then establishes support for the VNA if there are at least three reads spanning each the beginning of the VNA and its end in ref+. These two judgements are then used in the straightforward manner to classify the sample to be heterozygous, homozygous for the VNA, or homozygous for the hg18 allele. Some alleles can be classified as neither, if there is no evidence in hg18 and in ref+ (these alleles are then excluded from the analysis in the respective individual). The VNA frequency (VNAf) of an individual is the percentage of alleles at the validated sites that are those of Venter.

Unlike Delly, the validation classifier has a priori knowledge of the insertion or deletion sites and access to both the hg18 and ref+ alignments. This allows it to scrutinize the locus with single nucleotide resolution, so we consider it more reliable than Delly’s approach, which is oblivious to the differences between the two reference genomes. Additionally, it has access to higher coverage data. The classifier will nevertheless misclassify some alleles; however, it is not biased towards ref+ or hg18, so any potential misclassifications do not skew the results of our analysis.

## Acknowledgements

We would like to acknowledge Thanh Le for helping with alignment, Tobias Marschall and Alexander Schönhuth for initial discussions and providing us with VNA annotations, and Yao Wang for help with the gene analysis.

## Supporting information

**S1 Fig:** Venter Novel Alleles locations. We show the location of the VNAs and the genes they overlap. The figure is generating using the PhenoGram software [15].

**S2 Fig:** Proportion of validated VNA sites that have a VNA allele, per individual, segregated by population (as judged by the validation classifier).

**S1 Table:** Description of dataset.

**S2 Table:** VNA annotations and presence in samples. This spreadsheet contains information about each VNA and its status in each individual. The columns indicate the location of the VNA insertion in hg18, the sequence of the VNA, the gene (if any) which it overlaps, a column for each of the 16 individuals indicating its presence/absence as determined by the ref+ pipeline, a column for its status as indicated by the GRC pipeline, and a column for the status as determined by our validation classifier.

**S3 Table:** Pipeline accuracies. This table shows the accuracy of our ref+ and GRC pipelines for each of the individuals. For each sample, the validated sites are those for which our validation classifier finds evidence for at least one of the alleles (GRC or VNA).

**S4 Table:** Validation dataset

